# Chronic intermittent nicotine delivery with lung alveolar region-targeted aerosol technology produces circadian blood pharmacokinetics in rats resembling human smokers

**DOI:** 10.1101/195123

**Authors:** Xuesi M. Shao, Siyu Liu, Eon S. Lee, David Fung, Hua Pei, Jing Liang, Ross Mudgway, Jingxi Zhang, Jack L. Feldman, Yifang Zhu, Stan Louie, Xinmin S Xie

## Abstract

**Introduction:** Cigarette smoke is an aerosol containing microparticles that carry nicotine into lung alveolar region where nicotine is rapidly absorbed into circulation. Nicotine exposure in smokers is a chronic intermittent process, with intake during wakefulness and abstinence during sleep resulting in circadian fluctuation of blood nicotine levels. Here we present a smoking-relevant nicotine exposure device and rodent model.

**Methods:** We developed a computer controlled integrated platform where freely moving rodents can be exposed to episodic nicotine aerosol on an investigator-designed schedule. Rats were exposed to nicotine aerosol once every half hr in the dark phase of 12/12-hr dark/light cycles for 10 days. Plasma nicotine and its metabolite cotinine levels were determined with a LC-MS/MS method.

**Results:** We characterized the aerosol in the breathing zone of the rodent exposure chamber. The droplet size distribution was within the respirable diameter range. The system can generate a wide range of nicotine concentrations in air that meet a variety of experimental needs. We optimized the parameters of aerosol generation and exposure: plasma nicotine and cotinine concentrations reached 30-35 ng/ml and 190-240 ng/ml, respectively. The nicotine levels and circadian patterns resembled the pharmacokinetic pattern of human smokers.

**Conclusions:** We developed an aerosol system that can produce chronic intermittent nicotine exposure in unanesthetized and unrestrained rodents with route of administration and circadian blood pharmacokinetics resembling human smokers. This methodology is a novel tool for studies of behavior, pharmacology and toxicology of chronic nicotine exposure, nicotine addiction, tobacco-related diseases, teratogenicity, and for discovery of therapeutics.

**Implications:** We developed a method and an alveolar region-targeted aerosol system that provides chronic intermittent nicotine exposure in rodents. The method produces clinically relevant animal models with the route of administration and circadian pharmacokinetics resembling human smokers. This method is a novel tool for understanding the health effects of chronic nicotine exposures such as with tobacco cigarettes, E-cigarettes and other tobacco products, for studies of pharmacology, toxicology, nicotine addiction, tobacco-related diseases, and for discovery of medications.

## Introduction

Nicotine in tobacco products is highly addictive with significant adverse health consequences. Once smokers become addicted (approximately 1,200 per day in the United States), they have great difficulty quitting ^1^. In addition to conventional cigarettes, cigars and water pipes, devices for inhalation of nicotine such as non-combustion inhaler type of tobacco products^2^, the heat-not-burn tobacco heating systems^3^ and electronic nicotine delivery system (ENDS)/E-cigarette (E-cig)^4^ have emerged and become increasingly popular. Aerosols and vapors from these products are typically complex mixtures, but nicotine is the active addictive constituent shared by nearly all tobacco products. Therefore, understanding the health impacts of inhaled nicotine is critical.

Tobacco smoke is an aerosol containing microparticles in the aerodynamic size of 0.1–1.9 μm. Mass median aerodynamic diameter (MMAD) is 0.9-1 μm ^5,6^. When inhaled, particle size distribution determines the deposition behavior and deposition regions in the respiratory tract, including the upper airways, trachea, bronchi, bronchioles and alveolar regions^7^. Particles in the size range of cigarette smoke carry nicotine and other chemicals into the lungs, and are deposited in the alveolar regions where nicotine is quickly absorbed into pulmonary circulation. E-cigs also produce an aerosol^8-10^, not a simple nicotine vapor as incorrectly claimed by the E-cig industry. The characteristics of E-cig aerosol such as particle concentrations and size distributions have been measured and shown to be comparable to those of tobacco cigarette smoke^11-13^.

Nicotine addiction is a neurobiological adaptation to chronic nicotine exposure. Development of addiction depends on the forms of delivery, amount of nicotine delivered, rate of adsorption and nicotine pharmacokinetics (PK)^14,15^. Episodic smoking of cigarettes or E-cigs has distinct PK and activates nicotinic acetylcholine receptors (nAChR), followed by a cycle of desensitization/resensitization^16^. Nicotine exposure for smokers is a chronic intermittent process with episodic smoking during wakefulness and abstinence during sleep resulting in circadian fluctuations of blood nicotine levels. The episodic and circadian dynamics of nicotine exposure present a significant challenge to understand its long term effects; studies using chronic exposure, such as with osmotic pumps that deliver continuous amounts of nicotine, likely miss critical aspects of the effects of episodic nicotine, such as related to activation and desensitization/resensitization of receptors, as well as any circadian effects. The lack of smoking-relevant methods for nicotine delivery to animals and lack of clinically relevant animal models are major barriers of the research field.

We developed a non-invasive method with alveolar region-targeted aerosol technology for delivering nicotine to restrained rodents breathing through a nose-only chamber that produces blood PK resembling smoking a cigarette in humans^17^. Here, we present an integrated platform with computer control for unanesthetized and unrestrained, i.e., freely moving, rodents (rats or mice) that enables scheduled episodic nicotine aerosol deliveries, including circadian variability in exposure resembling that of human smokers and E-cig users.

## Methods

### Animals

All animal use procedures were in accordance with the National Institutes of Health (United States) Guide for the Care and Use of Laboratory Animals and were approved by the University of California, Los Angeles Institutional Animal Care and Use Committee. Efforts were made to minimize the number of animals used and their pain and suffering. Male Sprague-Dawley rats of 8 to 11-week-old (body weight 275–370 g) were used in this study. They were housed in the vivarium under a 12-hr light/dark cycle and had *ad libitum* access to food and water.

### Nicotine Aerosol Generation and Exposure System

The major component of the nicotine aerosol generation device is a 3-jet Collison nebulizer (BGI/Mesa Labs, Butler, NJ 07405, USA) that can consistently generate nicotine aerosol emissions having specific size distribution profiles. The system also contains an air pressure gauge (Ashcroft@ filled gauge, 0–100 psi. Cole-Parmer, Vernon Hills, IL, USA), two air flowmeters with valves (150-mm Direct Reading, 23 L/min, Cole-Parmer) for monitoring and regulating the inlet airflow rate and pressure applied to the nebulizer. The pressurized air was supplied from the air source of the laboratory facilities. The air pressure applied to the nebulizer for generation of the aerosol determines the flow, the droplet size distribution, and the concentration of the output aerosol. The air pressure and flow rate were well controlled and continuously monitored during the nicotine aerosol exposure experiments. The outlet of the nebulizer was opened to a sealed exposure chamber i.e., a plastic cylindrical tube of 19.7 cm inner diameter (ID) and 25.4 cm long with a platform that holds rodents. The exposure chamber mimicked a home-cage like environment where the rat can move freely during a long-term study. The volume for the rat free-moving space of the cylindrical chamber was around 6.7 L.

### Determinations of the aerosol droplet size distribution and mass concentration in air in the rat exposure chamber

The nicotine aerosol generation and exposure system was set up in a fume hood at temperature of 22 ± 1°C. Nicotine (freebase) was dissolved in NaCl solution or water for an osmolality ∼300 mOsm/kg. An Aerodynamic Particle Sizer (APS; model 3321, TSI Inc. Shoreview, MN, USA) was used to measure the droplet size distribution of nicotine aerosol generated from nicotine solution at different concentrations in the jar of the nebulizer. Total sampling airflow rate of APS was 5 L/min.

The sampling tube was located around the center above the platform where the rat is expected to stay on.

Mass concentrations of the nicotine aerosol were measured with a one-stage cascade impactor (SKC Inc. Eighty Four, PA, USA). Before the assay, the filters for the impactor were balanced in a temperature (22 °C) and humidity (49%) controlled environment where the electronic balance (Mettler-Toledo MX5 Microbalance, Mettler-Toledo Inc. Columbus, OH, USA) for weighing the filters was located. The filter weight difference between pre- and post-nicotine aerosol collection is the mass collected in a preset duration. Gravimetric filter sampling was conducted at sampling airflow rate of 2 L/min for 5 min in each experimental scenario.

### Nicotine aerosol exposure and blood sample collection

For blood sample collection, femoral artery catheterized rats were ordered from Harlan/Envigo Co (Indianapolis, IN, US).

For exposure accuracy, only one rat was put into each sealed exposure chamber. The device was programed to generate nicotine aerosol every half hour, for a total of 24 times in every 12-hour dark phase of 12/12-hour dark/light circadian cycles for 10 days. When the nicotine aerosol delivery pause (the time intervals between aerosol deliveries), fresh air from the pressurized air source was supplied to the chamber at a rate of 60 air change per hr (AC/h) adjusted with one of the two flow meter. 30-35 ml nicotine solution was put in the nebulizer jar and the solution was replaced with fresh one every day. Rats were returned to their home cage where no nicotine aerosol was delivered during the 12-hr light phrase.

For blood collection, the rat was put into a rat holder (Model #: CHT-250, CH Technologies Inc. Westwood, NJ, USA). Blood samples (equivalent to 0.1 ml plasma) were collected at a series of time points (see Results section) on day 3 and day 10. The blood was collected from the opening of the exteriorization tubing of the catheter into a 0.6-ml capillary blood collection tube containing ethylenediaminetetraacetic acid-K2 (Terumo Medical Co, Elkton, MD, USA). Samples were keep in ice and centrifuged within 15 min of collection at 1,000 g (gravity) for 12 min. The amount of each blood sample was around 200 μL and 100 μL plasma can be obtained. Supernatant plasma was transferred to labeled ice-cold eppendorf tubes and store below −20 °C. The plasma samples were sent to Louie laboratory at the School of Pharmacy, University of Southern California, Los Angeles, CA, USA for measurements of nicotine and cotinine.

### Measurements of plasma nicotine and cotinine concentrations

Plasma samples containing nicotine and cotinine were extracted using solid phase extraction method. In brief, 50 μL of internal standard (Nicotine-d3 10 μg/mL) was added to 50 μL plasma. Then 600 μL of 10% trichloroacetic acid was added to acidify the plasma sample. After centrifuging at 13,000 rpm for 5 min, supernatant was loaded onto pre-conditioned MCX SPE cartridge (Part # 186000252, Waters, USA). Cartridge was washed using 0.5 mL of 2% formic acid followed by 0.5 mL of methanol, and then analytes were eluted by 2X0.5 mL of methanol with 5% ammonium hydroxide. The eluted analytes were neutralize using 10 μL of 0.02 N HCl, and the samples were evaporated using a steady stream of dried and filtered nitrogen gas. Dried samples were reconstituted with 50 μL of mobile phase buffer and 40 μL was injected to LC-MS for analysis.

The concentrations of nicotine and its metabolite cotinine were quantified using a validated Liquid chromatography-tandem mass spectrometry (LC-MS/MS). The LC-MS/MS system consists of an API4000 triple quadrapole mass spectrometer (AB Sciex, USA) linked to Shimadzu LC-20AD HPLC (Shimadzu, Japan). All the analytes were detected by using multiple reaction monitoring in positive mode, where Q1→Q3 is 163.2→117.1 for nicotine, 177.2→80.1 for cotinine, and 167.2→121 for nicotine-d3 (internal standard), respectively. Analytes were separated using a Hypersil GOLD C18 column with parameters of 50 × 2.1 mm × 5 micron (Part #. 25005-052130, Thermo Scientific, USA). The mobile phase for the LC method included two components, water with 0.5% formic acid (Component A) and acetonitrile with 0.5% formic acid (Component B). A gradient program was used, where component B was started from 5% and increased gradually to 95% in 1 minute and held for another 3 minutes.

### Data Analysis

Plasma nicotine and cotinine levels were analyzed with two-way repeated-measures ANOVA using SigmaStat (Systat Software Inc. San Jose, CA 95131). Post-hoc multiple comparison based on Holm-Sidak’s method was used to determine significance levels between groups when needed.

### Chemicals

(s)-(-)-nicotine freebase (liquid, 99%) was purchased from Alfa Aesar Co (Tewksbury, MA, USA).

## Results

### Nicotine aerosol characteristics in the breathing zone of the rat free-moving exposure chamber

Generation of aerosol from a liquid depends on physical properties e.g., viscosity, vapor pressure and surface tension of the liquid. Once generated, the size distribution and concentration of aerosol change over time because of dynamic transformation process involving evaporation, coagulation and condensation^18^. Therefore, to deliver drugs to the body through the lungs in aerosol form, one needs to establish and verify specific procedures for different drugs based on the physical properties of the drugs and the solvents. In addition, characteristics of the nicotine aerosol, such as droplet size distribution and concentrations in air need to be measured at the breathing zone of the exposure chambers to obtain the information about the aerosol inhaled by the experimental animals. Respirable diameter is defined as the aerodynamic diameter of aerosol particles capable of reaching the gas exchange region in the lungs for the organism under study ^19^. For inhalation toxicology studies, a respirable droplet size range of MMAD between 1–4 μm is recommended^19^ (Also refer to ^20^)

The aerosol droplet size distribution in the breathing zones of the exposure chamber that allows the rat to move freely is shown in Figure 1. When the nicotine concentrations in the nebulizer container were low (0.01%-0.1%), the droplet size distributions were close to a log-normal with a slight elevation in the range of large droplets. Varying the air pressure applied to the nebulizer in the range of 20-60 psi had little effect on the distribution (Fig. 1A). In contrast, high concentrations of nicotine (10%) resulted in a bimodal size distributions, probably due to higher viscosity of the solutions. The numbers of larger size droplets decreased with increase in air pressure (Fig. 1B). Aerosol mass size distributions were summarized as MMAD, which substantially increased as nicotine concentrations in the nebulizer increased (Table 1). With solutions of low nicotine concentrations in the nebulizer (from 0.01%-1%), MMAD slightly increased as air pressure increased in the range of 20-60 psi. With high nicotine concentration (10%) in the nebulizer, MMAD slightly decreased as air pressure increased (Table 1). The MMAD values in our experimental conditions are within the respirable size range recommended by US EPA^19,20^. These data are consistent with those of the output data documented by the manufacture, BGI Inc. (http://bgi.mesalabs.com/collison-nebulizer/).

**Figure 1.**
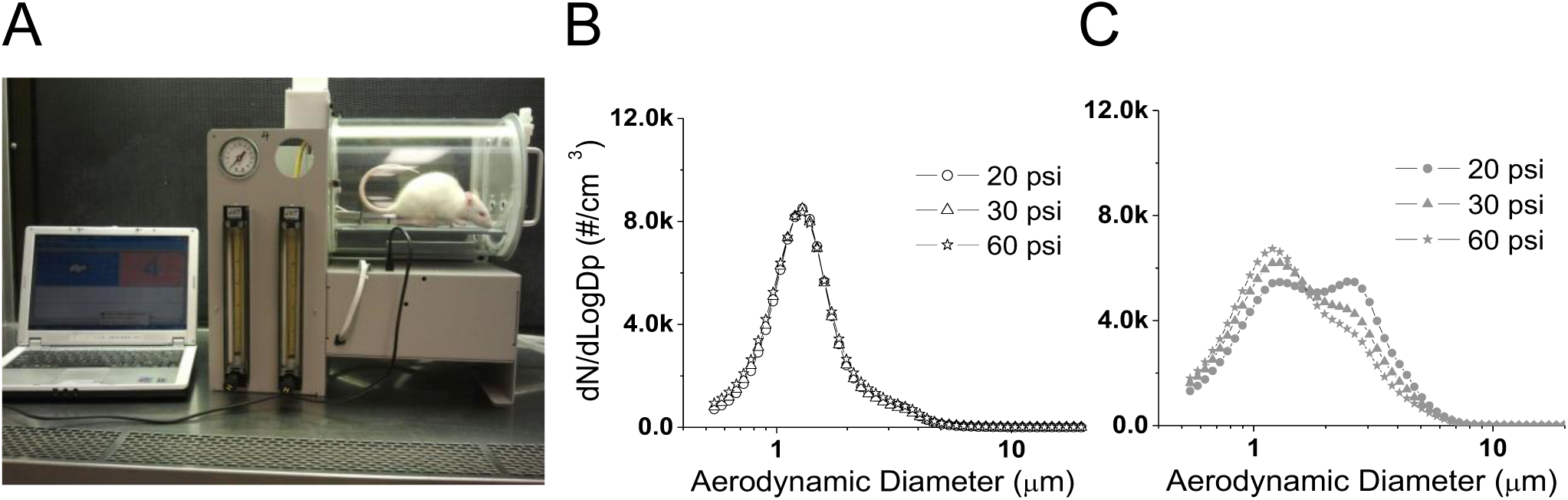
The nicotine aerosol droplet size distribution in the breathing zone of the rat free-moving exposure chamber. (A) The aerosol generation and rodent exposure system. The nicotine concentration in the 3-jet Collison nebulizer container was (B) 0.1%; (C) 10%. Air pressure in the inlet of the nebulizer was 20, 30 or 60 psi. The data are averages of 60 samples of 1 second during a total 1 min aerosol sampling time using an Aerodynamic Particle Sizer^^®^^ (APS). The lower detection limit was 0.52 μm in aerodynamic diameter and sampling airflow rate was 5 L/min.

**Table 1.** Nicotine aerosol droplet size, aerosol mass concentrations (conc) and nicotine conc in air in the breathing zone of the free-moving chamber for rats

| Nicotine conc in nebulizer | Air pressure (psi) | MMAD (μm) | GSD | Aerosol mass conc in air (mg/L) | Nicotine conc in air (mg/L) |
| --- | --- | --- | --- | --- | --- |
| 0.01% | 20 | 2.78 | 1.72 | 1.45 | 0.000145 |
| 0.01% | 30 | 2.88 | 1.72 | 1.74 | 0.000174 |
| 0.01% | 60 | 2.90 | 1.72 | 2.3 | 0.00023 |
| 0.1% | 20 | 2.83 | 1.76 | 1.91 | 0.00191 |
| 0.1% | 30 | 2.86 | 1.78 | 1.96 | 0.00196 |
| 0.1% | 60 | 3.07 | 1.76 | 2.16 | 0.00216 |
| 1% | 20 | 3.15 | 1.67 | 1.62 | 0.0162 |
| 1% | 30 | 3.19 | 1.72 | 2.02 | 0.0202 |
| 1% | 60 | 3.30 | 1.71 | 2.12 | 0.0212 |
| 10% | 20 | 3.85 | 1.53 | 1.8 | 0.18 |
| 10% | 30 | 3.82 | 1.60 | 1.97 | 0.197 |
| 10% | 60 | 3.75 | 1.64 | 2.25 | 0.225 |

MMAD: mass median aerodynamic diameter. GSD: geometric standard deviation. psi: pounds per square inch. Conc: concentration. Aerosol was sampled using an Aerodynamic Particle Sizer® (APS). The APS sampling time was 1 min at a flow of 5 L/min.
Mass concentrations of the nicotine aerosol were measured with a one-stage cascade impactor.
Each sample was collected for 5 min in a flow of 2 L/min after 1/2 min for equilibrium of aerosol in the rat chamber. Nicotine concentrations in air (Cn) is estimated: $C_n = C_m * C_s$
Where Cs is the nicotine concentration in the solution in the nebulizer and Cm is the aerosol mass concentration in air.

These data suggest that the droplet size distribution depends on the physical properties of the solution and the nebulizer air pressure, as solutions with higher nicotine concentrations are higher in viscosity and lower in vapor pressure. Under a wide range of experimental conditions, rodents in the free-moving chamber were exposed to nicotine aerosol with respirable droplet size that can get into the alveolar region of the lungs.

Next we determined the nicotine aerosol concentrations in air in the breathing zone of the free-moving rodent chamber. As shown in Table 1, the aerosol mass concentration in air increased with increase of air pressure applied to the inlet of the nebulizer and with increase of nicotine concentration in the solution in the nebulizer container. The nicotine concentration in air (Cn) depends on the aerosol mass concentration in air (Cm) and nicotine concentration in the solution (Cs) in the nebulizer (Table 1 legend).

The droplet size distributions and mass concentrations were similar at different location in the cylindrical exposure chamber except the very corners of the cylinder.

These results indicate that our system can deliver a wide range of amounts of nicotine quickly into the circulation of rodents through inhalation route in a free-moving cage. The nicotine dose can be readily controlled by altering the nicotine solution concentrations in the nebulizer or other aerosol generation parameters, i.e., air pressure in the nebulizer. Our data can serve as a guide for choosing parameters in nicotine aerosol inhalation experiments for researchers to develop rodent models simulating human tobacco and E-cig smoking.

### Chronic intermittent nicotine aerosol (CINA) inhalation exposure in rats produces circadian blood PK resembling human smokers

The parameters of aerosol generation including nicotine concentrations in the solution in the nebulizer, the air pressure entering the nebulizer, the duration and interval of aerosol deliveries were adjusted and optimized based on measurements of nicotine and cotinine levels in the plasma of exposed rats. A solution of 0.1% nicotine in 0.9% NaCl (adjusted to pH 8.0 with HCl) was put in the container of the nebulizer. We delivered nicotine aerosol in a schedule of once every 30 min for a total of 24 deliveries in every 12-hr dark phase of 12/12-hr dark/light circadian cycles for 10 days. The duration of nicotine aerosol delivery was 1 min except the 1^st^ daily aerosol delivery which was 2 min in order to boost the nicotine level from the lowest level at the beginning of the dark phase. We previously reported that acute nicotine aerosol inhalation produces a quick rise in arterial blood nicotine levels followed by a slower decline. The arterial nicotine levels decrease to a level close to that of venous blood in ∼30 min^17^.

On day 3 and day 10, we collected blood samples every two hrs by the end of the 29 min intervals between aerosol deliveries when arterial nicotine level is similar to that of venous blood during the 12-hr dark phase, and every 4 hrs during the 12-hr light phase (when there was no nicotine aerosol exposure). As shown in Fig. 2, plasma nicotine concentrations gradually increased during the 12-hr dark phase, reaching a peak of 30-35 ng/ml in 8-12 hr from the start of the daily nicotine aerosol deliveries, with a slower rising of cotinine levels reaching 190-240 ng/ml. Nicotine levels gradually decreased to 5-10 ng/ml and cotinine to 70-100 ng/ml during the light phase. There was no significant difference between day 3 and day 10. Summary nicotine PK parameters are listed in Table 2. Area under the concentration-time curve (AUC_0-24hrs_) indicates daily total systemic nicotine exposure in our CINA inhalation exposure conditions.

**Figure 2.**
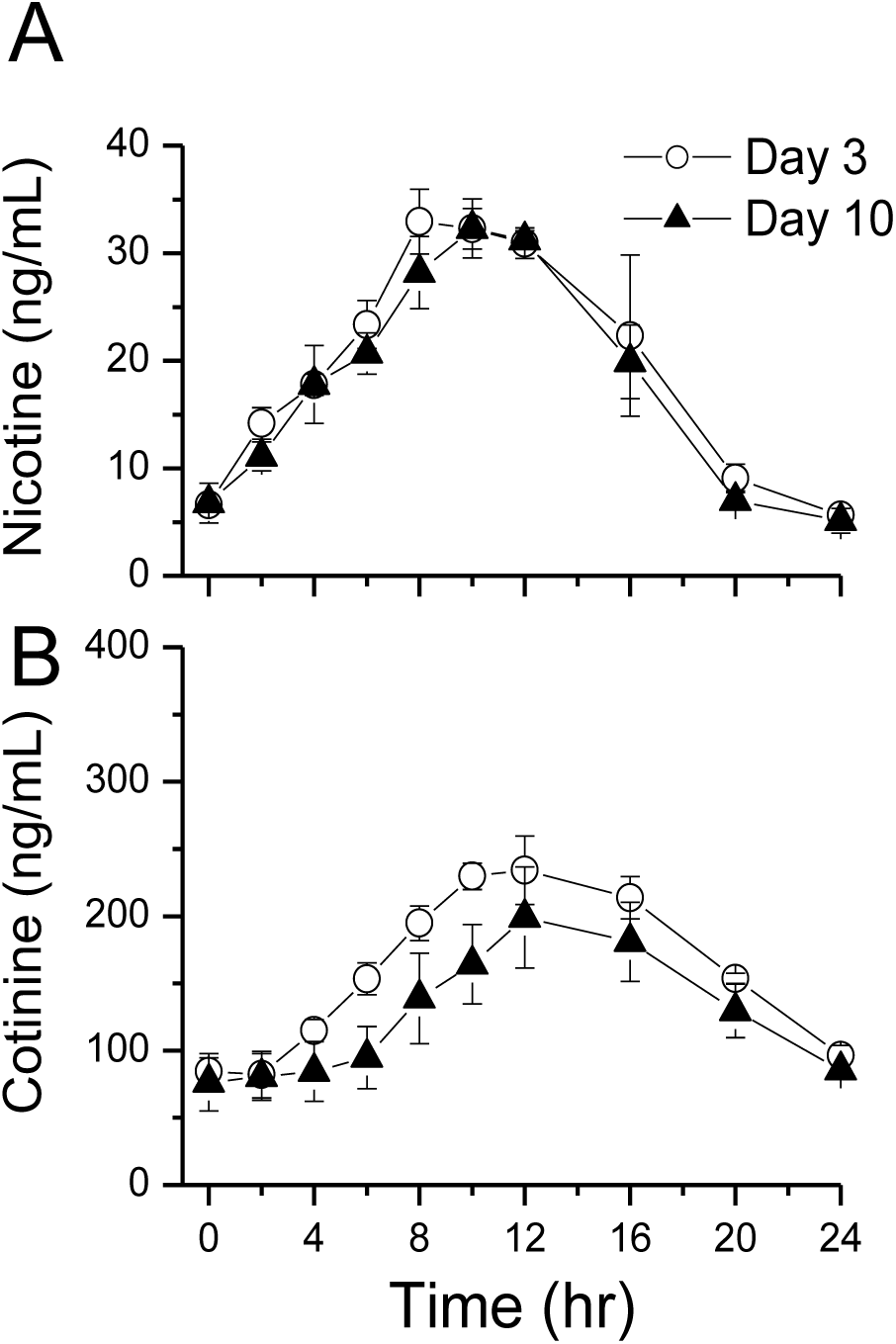
Chronic intermittent nicotine aerosol (CINA) delivery to rats produces circadian blood pharmacokinetics resembling that of human smokers. The air pressure for aerosol generation was 30 psi. At this pressure, the air flow in the rodent exposure chamber was 7.1 LPM (Volume under standard conditions, calculated based on pressure of 30 psi). Rats in the free-moving chamber were exposed to nicotine aerosol once every 30 min for a total of 24 deliveries in the 12-hr dark phase of 12/12-hr dark/light circadian cycles for 10 days. The duration of each nicotine aerosol delivery was 1 min except the 1^st^ one of the daily aerosol deliveries which was 2 min. Blood samples were collected from the femoral artery catheter on day 3 and day 10. (A) plasma nicotine levels (mean ± SE). (B) cotinine levels. Data at time point 0 are pre-nicotine control levels before the first daily nicotine aerosol delivery. There is no statistically significant difference between day 3 and day 10 (Two-way repeated-measures ANOVA, p = 0.39 for nicotine and p = 0.3 for cotinine. n = 4).

**Table 2.** Nicotine pharmacokinetic parameters with CINA exposure

|  | C<sub>max</sub> (ng/ml) | T<sub>max</sub> (hrs) | Half-life (hrs) | AUC<sub>0-24 hrs</sub>(ng/ml) |
| --- | --- | --- | --- | --- |
| Day 3 | 38.8 ± 4.33 | 10.5 ± 3.79 | 4.6 ± 1.57 | 479.8 ± 51.84 |
| Day 10 | 35.3± 2.42 | 10 ± 1.63 | 4.5± 1.47 | 438.4 ± 56.47 |

Mean ± SD. C<sub>max</sub>: the peak plasma concentration. T<sub>max</sub>: time to reach C<sub>max</sub>. AUC<sub>0-24 hrs</sub>: Area under the concentration-time curve from 0 to 24 hrs estimated with linear trapezoidal method.

These data indicate that the circadian fluctuation patterns of blood nicotine and cotinine levels in our rat model are similar to the circadian blood (venous) PK of human smokers^21^, with an exception of lower nicotine and cotinine levels by the end of the light phase, which is consistent with the notion that nicotine metabolism is faster in rats compared to humans^22^. The circadian PK was stabilized from day 3 and consistent throughout the CINA treatment period of 10 days, suggesting a balance of nicotine intake vs. elimination and metabolism under our experimental conditions.

## Discussion

We developed a clinically relevant, non-invasive method and a novel aerosol system for CINA inhalation/exposure for rodents. 1) The particle size of the nicotine aerosol in the breathing zone of the exposure chamber is of respirable diameter^19^, indicating that nicotine aerosol deposits in the alveolar region to quickly enter the pulmonary circulation, just like tobacco and E-cig smoke. 2) A wide range of nicotine concentrations in air can be achieved in this system, therefore, the dose of inhaled nicotine can be readily controlled for variety of experimental rodent models for studies on, e.g., smoking a cigarette/E-cig, chronic smokers and inhalation toxicology. The nicotine concentration can be 3-4 orders of magnitude higher than that of the vapor inhalation method where nicotine concentration is 0.001 ± 0.00002 mg/L in air^17,23^. 3) The nicotine aerosol inhalation/exposure method is efficient and consistent. CINA inhalation during wakefulness produced circadian PK in rats resembling that of chronic smokers. Due to the small blood volume of rats that limits the numbers of blood samples, further studies are needed to simulate puff by puff PK. We previously reported that nicotine aerosol inhalation using a nose-only device achieved a quick rise in arterial blood nicotine levels followed by a slower decline, similar to the PK pattern observed in humans smoking a cigarette. In those conditions, the rat was restrained in a rat holder ^17^. Here, freely-moving rodents exposed to CINA achieve plasma nicotine kinetics and the plasma levels comparable to chronic human smokers. Nicotine can be administered with this aerosol inhalation/exposure method in vivo under a broad range of experimental conditions including freely behaving, sleeping, restrained, anesthetized, or artificially ventilated rodents. The devices and method represents a powerful tool for studies of nicotine addiction, nicotine toxicology, tobacco-related diseases affecting, for example, the CNS, the lungs, the cardiovascular system, and, if applied to pregnant rodents, nicotine teratogenicity and for discovery of pharmacological therapeutics.

For animal models for studies of drugs of abuse, the route of exposure and the PK or target tissue concentrations of the drug should be comparable between animals and humans. Conventional chronic nicotine exposures for animals have significant limitations. (1) Oral nicotine administration through drinking water^24^. Nicotine absorption is slow compared to smoking. Blood nicotine levels achieved by oral intake are affected by first-pass liver metabolism. (2) Subcutaneous osmotic mini-pumps^25-27^. The nicotine delivery is slow and cannot simulate episodic exposure as smoking cigarettes in humans. (3) Intravenous nicotine self-administration through a chronically implanted catheter. This method is invasive and maintaining the i.v. catheter patency for a long period of time is technically challenging^28,29^. (4) Nicotine vapor inhalation method^23,30^. Nicotine concentrations in vapor form in air is low, not comparable to cigarette or E-cig smoke. Furthermore, the amount of nicotine that enters circulation is limited because nicotine vapor mostly deposits and is absorbed in nasal and buccal mucosa, while very little deposits in the lungs^31,32^. Animals have to be continuously exposed to nicotine vapor for hours to produce blood nicotine concentrations comparable to that of human smokers^23^. In contrast, with nicotine aerosol inhalation/exposure method as tobacco cigarette or E-cig smoking, aerosol deposits in the alveolar region of the lungs, peak levels of nicotine are higher and lag time between smoking and entry into the brain shorter. Rapid onset of effects in the brain provides an optimal reinforcement schedule for the development of nicotine dependence^33^. In addition, aerosol inhalation is a powerful method for studying clinically relevant impacts of nicotine on the pulmonary system, hemodynamics and cardiovascular system. Compared to nicotine entering the body through oral or dermal routes, inhaled nicotine has distinct PK and undergoes less metabolic processing before encountering components of the lungs and cardiovascular systems.

The dose of nicotine administered through inhalation to rats can be roughly estimated. Minute ventilation/100 g body weight of Sprague-Dawley rats is 28.17 ± 1.37 ml^34^.

Inhaled dose = nicotine concentration in air * minute ventilation * exposure time^35^. For example, when the nicotine concentration in the nebulizer is 0.1% and air pressure is 30 psi, we have nicotine concentration in air = 0.00196 mg/L (Table 1). Therefore

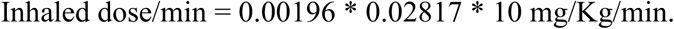

We showed, in a previous study, that a single acute nicotine aerosol inhalation exposure can produce arterial and venous PKs resembling that of human smoking a cigarette ^17^. However, PKs produced by CINA exposure are expected to be substantially different. Nicotine apparent volume of distribution in rats is very high as 4.7-5.7 L/Kg^36^. In humans, this value, on average, in steady state is 2.2-3.3^22^. The decline phrase after a single nicotine aerosol exposure is determined primarily by distribution half-life while in CINA exposure, the decline phrases are determined by elimination half-life and rates of distribution out of storage tissues ^33^. It suggests that much higher doses are needed in acute single nicotine aerosol exposure experiments in rats in order to achieve comparable nicotine PK in humans, since blood nicotine quickly distributes to body tissues. In contrast, low doses are needed in CINA exposure conditions. The data in our previous study on acute nicotine inhalation/exposure conditions ^17^ and data obtained in CINA experiments in this study are consistent with this hypothesis.

One limitation of freely-moving whole body nicotine aerosol exposure for rodents is a possible dermal and/or oral exposure, e.g., by licking their fur where the aerosol could have condensed. We estimate that this contribution to the blood nicotine PK is insignificant as we obtained a circadian PK pattern similar to human smokers and the blood nicotine was very low toward the end of the 12-hr light phrase.

